# Role of Mce proteins in *Mycobacterium avium paratuberculosis* infection

**DOI:** 10.1101/2023.06.06.543821

**Authors:** Rose Blake, Kirsty Jensen, Omar Alfituri, Neil Mabbott, Jayne Hope, Jo Stevens

## Abstract

*Mycobacterium avium* subspecies *paratuberculosis* (MAP) is the causative agent of Johne’s Disease, a chronic granulomatous enteritis of ruminants. MAP establishes an infection in the host via the small intestine. This requires the bacteria to adhere to and be internalised by cells of the intestinal tract. The effector molecules expressed by MAP for this remain to be fully identified and understood. The mammalian cell entry (Mce) proteins play an essential role for other Mycobacterial species to facilitate the attachment and invasion of host epithelial cells. Here, we have expressed Mce1A, Mce1D, Mce3C and Mce4A proteins derived from MAP on the surface of a non-invasive *E. coli* host to characterise their role in the initial interaction between MAP and the host. To this end, *mce1A* was found to significantly increase the ability of the *E. coli* to attach and survive intracellularly in THP-1 cells, and *mce1D* was found to significantly increase attachment and invasion of MDBK cells. Both genes were implicated in the increased ability of *E. coli* to infect 3D bovine basal-out enteroids. Together, these results have identified two effector molecules used by MAP with a degree of cell-type specificity.

## Introduction

*Mycobacterium avium* subspecies *paratuberculosis* (MAP) is a Gram positive, facultative intracellular rod-shaped bacteria which is the causative agent of Johne’s Disease (JD). JD is a chronic gastric enteritis which affects ruminants across the world. Animals are typically infected within the first 6 months of life via the ingestion of contaminated milk and feed (Larsen et al., 1975). Other routes of infection include horizontal transfer from the environment and wildlife reservoirs such as rabbits, foxes and stoats (Corn et al., 2005; Judge et al., 2006).

After ingestion, MAP will travel to the small intestine of the animal and will target specific cells in the intestinal wall to traverse to establish an infection. M cells are known to be a target of several enteric pathogens, such as *Salmonella*, as these are antigen targeting cells which can uptake pathogens in the lumen of the intestine and present the pathogen to the underlying Peyer’s Patches (Wang et al., 2015). Immune cells such as macrophages and dendritic cells reside within the Peyer’s Patch to phagocytose pathogens which are presented to it. This enables the host to generate an appropriate immune response to eliminate the infection. MAP, like other Mycobacteria, takes advantage of this process to be phagocytosed by macrophages where it can survive and replicate (Arsenault et al., 2014). This initiates a chronic inflammatory response which may lead to the formation of granulomas in the intestinal lining, and clinical disease in 10% of infected animals (Arsenault et al., 2014; Lombard, 2011).

It may take 2-5 years from the initial infection before the animal may show clinical disease, which typically presents as chronic diarrhoea and emaciation of the animal. Within the subclinical period, MAP can be shed intermittently in the faeces and serve to infect other animals within the herd, yet remain below the level of detection for many diagnostic tests (Nielsen & Toft, 2008). Due to the limited diagnostic tests currently available for MAP, it is critical to identify factors expressed by MAP which may aid its infection of the host and may serve as therapeutic targets.

Currently, the initial interaction between MAP and the host remains largely unknown. While MAP has been shown to target M cells in a murine model using the FAP expressed on its surface (Secott et al., 2004), some researchers question the importance of this cell type in establishing a MAP infection. In juvenile cattle, the cell types of the small intestine are different than that of fully mature cattle. There are fewer M cells as these have not been stimulated to be matured yet. In addition, the FAP adhesion observed to aid infection of M cells did not aid the infection of enterocytes in the same study. This has led to the conclusion that MAP may express other adhesins on its surface which aid the infection of other cell types of the intestine including enterocytes (Bannantine et al., 2003; Bermudez et al., 2010) and goblet cells (Schleig et al., 2005), and therefore may provide a critical route for MAP to establish an infection.

Mammalian cell entry (mce) operons are highly conserved between Mycobacterial species. Recombinant *E. coli* or latex beads expressing *mce1A, mce3C* and *mce4A* have been observed to confer increased attachment and invasion of mammalian cells for multiple Mycobacterial species including *M. tuberculosis* and *M. bovis, M. leprae* (Saini et al., 2008; Casali & Riley, 2007; Zhang et al., 2018). In addition, *mce1A* derived from *M. tuberculosis* has been shown to increase intracellular survival of the recombinant expression host (Arruda et al., 1993). Due to the highly homologous nature of these operons between Mycobacterial species, it was hypothesised that the *mce* genes in MAP would function similarly to increase attachment and invasion of MAP to host cells. This was supported by a transposon mutagenesis study which observed that mutating *mce1D* in MAP resulted in a significant reduction of MAP infection in MDBK cells (Alonso-Hearn et al., 2008).

In the present study we investigate the role of several *mce* genes expressed by MAP using a recombinant *E. coli* host. MDBK cells, THP-1 cells and bovine intestinal organoids (enteroids) were used to investigate cell tropism displayed by *mce1A, mce1D, mce3C* and *mce4A*.

## Materials and Methods

### Bacterial isolates and culture conditions

*E. coli* were cultured in LB broth containing the relevant antibiotics where appropriate at 37°C 180 rpm. MAP isolates were cultured in 7H9 medium (270 mL water; 1.41g Middlebrook 7H9 supplement; 333 μL glycerol; 333 μL Mycobactin J; 30 mL OADC supplement) at 37°C 100 rpm.

### MAP exposure to acid

Five mL of MAP K10 and MAP C49 in the log phase were transferred to 50 mL falcon tubes and centrifuged at 3220 *x g* for 10 minutes. The cultures were re-suspended in either 5 mL standard 7H9 medium or 5 mL of 7H9 medium made to pH 3.0 prior to autoclave. Cultures were then immediately centrifuged at 3220 *x g* for 10 minutes for RNA isolation to serve as a control, or cultured at 37°C 100 rpm for 2 hours. The cultures were then centrifuged at 3220 *x g* for 10 minutes for RNA isolation. All experiments were carried out in three biological replicates.

### RNA isolation from MAP

Immediately after pelleting, bacterial pellets were re-suspended in 1 mL Trizol reagent (Thermo Fisher Scientific) and homogenised using lysing matrix B beads (MP Biomedicals). Tubes were pulsed in a Fastprep machine at 6.0 speed for 30 seconds twice followed by 6.5 speed for 45 seconds, keeping samples on ice for 5 minutes between steps. The sample was centrifuged at 16,200 *x g* for 3 minutes and the supernatant aliquot into a screw-top micro-centrifuge tube. 200 μL chloroform was added to the suspension and the sample treated for RNA isolation as described by the manufacturer, with some adjustments. Briefly, after centrifugation with isopropanol only 50% of the supernatant was removed, and all subsequent centrifugation steps were performed at 12,000 x *g* for 20 minutes. RNA precipitation was performed with 75% ethanol which was centrifuged at 7500 x *g* for 10 minutes.

### RT-qPCR

cDNA was synthesised from the RNA using Agilent AffinityScript Multiple Temperature cDNA Synthesis Kit. The oligo(dT) primers supplied were used and the protocol followed according to the manufacturer’s instructions. No reverse transcriptase controls were used to confirm the absence of genomic DNA from the samples.

All qPCR experiments were performed using SYBR green Supermix (Quantabio, VWR international Ltd). The total reaction volume was 10 μL consisting of of 5 μL Supermix, 0.5 μL forward primer, 0.5 μL reverse primer, 1.5 μL nuclease free water and 2.5 μL template. Samples were loaded in triplicate into 96 well plates and no template controls were included to verify the absence of contamination. Oligonucleotides were designed using Primer3 (Koressaar & Remm, 2007; Untergasser et al., 2012) and Netprimer (Biosoft International) software and are outlined in Table 1 using the “conventional” PCR primers to generate PCR amplicons to act as the template for individual standard curves. The experimental cDNA was diluted 1:20 to generate template for the RT-qPCR reaction. The relative quantities of mRNA were calculated using the Pfaffl method (Pfaffl, 2001), using the geometric mean of the RT-qPCR results for the reference genes *GAPDH* and *1g2* (Granger et al., 2004; Pribylova et al., 2011) to calculate differences in the template RNA levels for standardisation of the Ct values for the genes of interest.

**Table 1.**
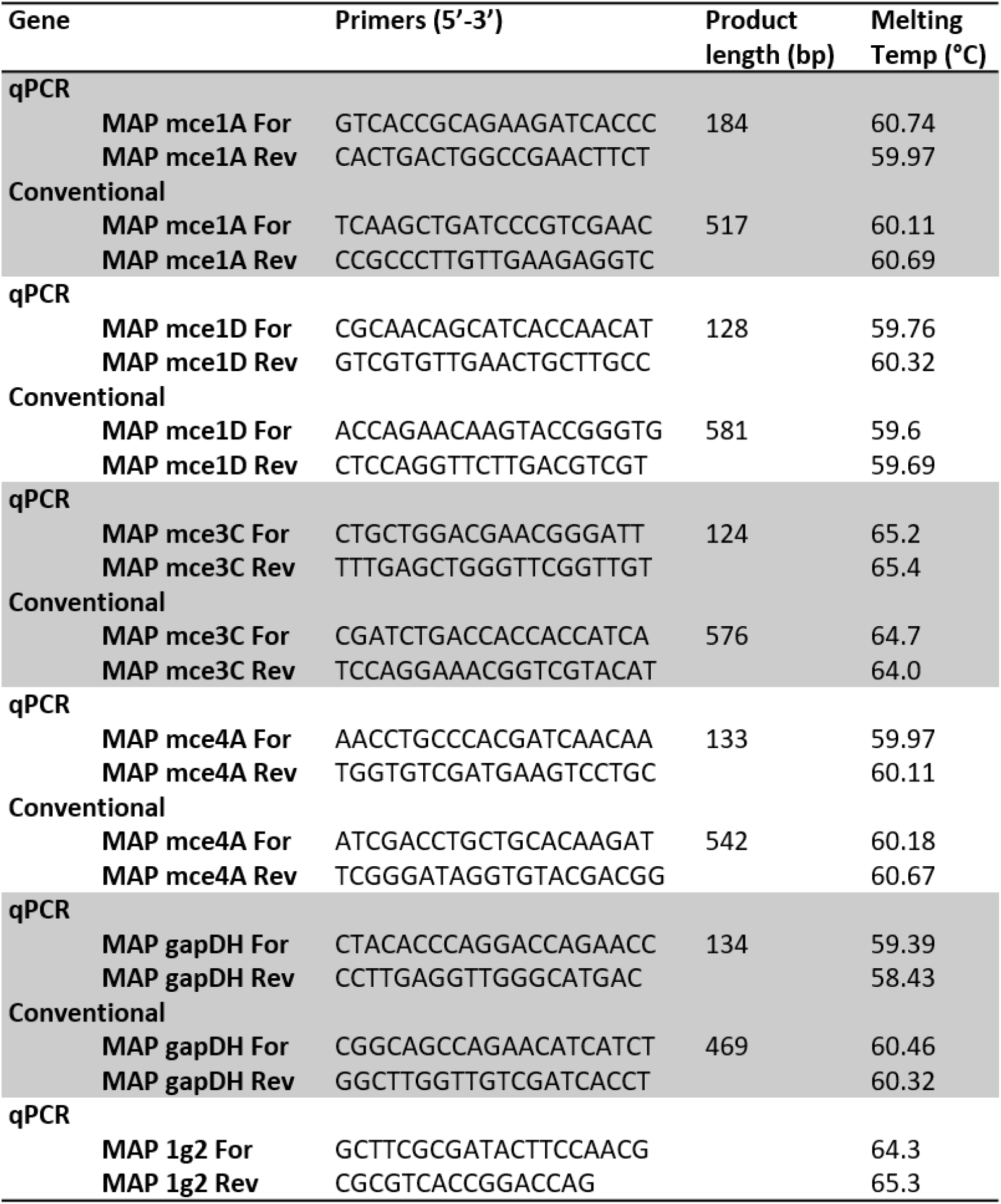
Primer pairs used to amplify and then quantify specified genes using qPCR.

### Cloning, expression and expression of Mce1A, Mce1D, Mce3C and Mce4A

Genomic DNA from *Mycobacterium tuberculosis* H37Rv was gifted by Dr. Jordan Mitchell to act as a positive control, and genomic DNA from MAP K10 was extracted using the Qiagen Blood and Tissue kit. Full length *mce1A* from *M. tuberculosis* (Rv0169), *mce1A* from MAP K10 (MAP3604), *mce1D* from MAP K10 (MAP3607), *mce3C* from MAP K10 (MAP2114c) and *mce4A* from MAP K10 (MAP0564) genes were amplified from genomic DNA by PCR using the primers listed in Table 2.

**Table 2.**
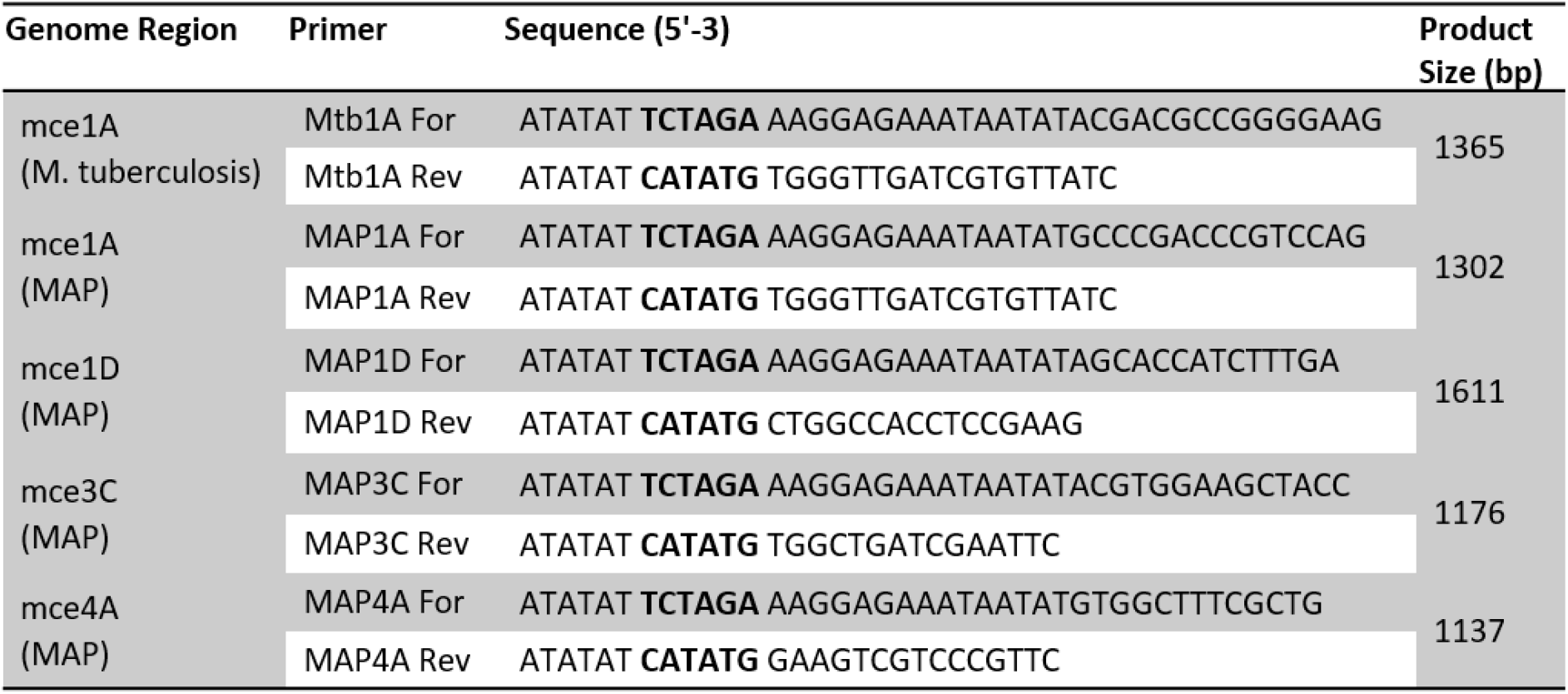
Primer pairs used to amplify the appropriate mce gene for subsequent cloning steps. Restriction enzyme sites are in bold

These genes will be referred to as *mtb1A, map1A, map1D, map3C* and *map4A* respectively henceforth. Each forward primer contained an *Nde*I restriction enzyme site and each reverse primer contained an *Xba*I restriction site.

PCRs were performed in 25-50 μL volumes using 10 ng template DNA, 1 μM oligonucleotide primers, 200 μM deoxynucleotides triphosphate, 0.04 U/mL Phusion High Fidelity Polymerase, 1x GC buffer, 6-10% DMSO and 5 mM MgCl_2_ depending on the primer pair used. 35 cycles of DNA amplification were performed. The samples were denatured at 98°C for c. 5 minutes, annealed at c. 52°C and extension at 72°C (allowing 30 seconds per 500 bp of DNA to be synthesised). The amplicons were cloned into pET21b(+) to generate the appropriate *mce* construct with a hexa histidine tag. Both DH5α and Rosetta 2 (DE3) strains of *E. coli* were transformed with the recombinant plasmids for expression studies. The presence of the inserts was confirmed by PCR, restriction enzyme digestion and sequencing.

Overnight cultures of *E. coli* Rosetta 2 (BL-21) (Novagen) containing the recombinant expression plasmids was diluted 1:10 in fresh LB broth containing ampicillin and chloramphenicol and cultured at 37°C 180 rpm to an optical density (OD) of 0.6 at 600 nm. For transcription induction, IPTG (isopropyl thio-b-D-galactoside, Sigma) to a final concentration of 0.1 mM was added to the culture and incubation was continued for 2 hours.

### Subcellular fractionation

Recombinant *E. coli* were cultured in 200 mL LB broth containing ampicillin and chloramphenicol at 37°C 180 rpm to an OD of 0.6 at 600 nm and then Mce protein expression was induced with 0.1 mM IPTG for a further 2 hours. Cells were harvested by centrifugation at 3000 x *g* for 10 minutes. The pellet was washed with 0.1 volume of TM buffer (20 mM Tris-HCl pH 7.0, 3 mM MgCl_2_) and repelleted at 3000 x *g* for 10 minutes and frozen at -80°C. The pellet was then suspended in 3 mL of 10 mM Tris-HCl pH 7.0, 25% sucrose (w/v). A cocktail of protease inhibitors (Sigma) was added at five mL per gram of wet pellet and lysozyme (SERVA) was added to 0.5 mg/mL (w/v) and incubated at 37°C for 20 minutes. MgCl_2_ was then added to a final concentration of 3 mM and incubated at 37°C for 20 minutes. One volume of 4% Triton-X100 was added and mixed for 4 minutes before freezing the solution at -80°C and subsequently thawed at 37°C and mixed for 1 minute. This freeze-thaw cycle was repeated for a second time before the supernatant was removed after centrifuging the sample at 7500 x *g* for 15 minutes and frozen at -80°C. The supernatant was thawed at RT and ultracentrifuged at 110,000 x *g* for 1 hour at 5°C to pellet the crude outer membranes.

The supernatant was stored and represents the cytoplasmic fraction of the bacteria; the pellet was resuspended in 0.3 mL TM by bath sonification for 5 minutes. One volume of 4% Triton-X100 was added and incubated on ice for 30 minutes before centrifuging at 7500 x *g* for 15 minutes. The supernatant was aliquot into a separate tube and ultra-centrifuged at 110,000 x *g* for 1 hour. The pelleted membranes were suspended in 0.4 mL TM by sonification and centrifuged at 7500 x *g* for 15 minutes. The supernatant containing the outer-membranes of the bacteria were removed and diluted 4-fold with 5 mM Tris-HCl pH 8.0. The outer membranes were pelleted by ultracentrifugation at 110,000 x *g* for 1 hour. The pellet was suspended using sonification in 0.6 mL 5 mM Tris-HCl pH 8.0 and RNase was added to 0.5 mg/mL (w/v). This was incubated at 37°C for 10 minutes and EDTA was added to 10 mM and incubated for a further 40 minutes. The sample was then adjusted to 50 mM Na_2_CO_3_ and 1 M NaCl and incubate on ice for 1 hour before incubation at 37°C for 15 minutes. The membranes were centrifuged at 110,000 x *g* for 1.25 hours and washed by sonification in 0.1 mL 100 mM Na_2_CO_3_ and 1 M NaCl. The sample was incubated on ice for 30 minutes and pelleted at 110,000 x *g* for 1 hour. The pellet was washed again by sonification in 0.1 mL 100 mM Na_2_CO_3_ and 1 M NaCl and re-pelleted as described.

The final fractions representing the cytoplasm and membrane, cytoplasm alone, and 2 membrane fractions at different wash periods were selected to analyse by western blot to detect the presence of the His-tagged Mce proteins.

### Detection of protein expression

To detect the C-terminal His-tagged recombinant Mce protein in the *E. coli*, anti-His monoclonal antibody (Raybiotech) and goat anti-rabbit IgG were used as primary and secondary antibodies (Supplementary Table 1). For the intracellular localisation of the Mce proteins in *E. coli*, Western blot was performed on samples containing the cytoplasmic and membranous fractions, the cytoplasmic fraction alone, and two membrane fractions at different wash periods.

### Invasion assays

Invasion of MDBK cells seeded at 1 × 10^5^ cells per well into a 24-well plate by Rosetta 2 (BL-21) transformed with pET21b(+)/*mtb1A*, pET21b(+)/*map1A*, pET21b(+)/*map1D*, pET21b(+)/*map3C*, pET21b(+)/*map4A* and empty vector pET21b(+) were assayed. *E. coli* were cultured overnight and diluted 1:10 in fresh medium containing ampicillin and chloramphenicol and incubated at 37°C 180 rpm to reach an OD of 0.6 at 600 nm and induced with 0.1 mM IPTG for 2 hours at 37°C. Following induction, the bacteria was normalised to an OD of 0.6 at 600 nm and centrifuged at 3000 x *g* for 10 minutes and re-suspended in mammalian cell culture medium. Prior to the infection, fresh medium replaced the mammalian cell culture medium. Recombinant *E. coli* cells were added to the monolayer at a multiplicity of infection (MOI) of 20:1 for MDBK cells and an MOI of 10:1 for THP-1 cells and incubated at 37°C for 1 hour. The cells were washed three times with PBS, and fresh cell culture medium was added to the infected cells for a further 1 or 5 hours for incubation at 37°C. For THP-1 invasion assays, the fresh medium contained 10 μg/mL gentamicin to kill extracellular bacteria.

### Enteroid generation and infection

3D basal-out bovine enteroids were maintained as described in (Blake et al., 2022). Briefly, enteroids were suspended in Matrigel domes and cultured with murine IntestiCult (STEMCELL) supplemented with 10 μM Y-27632, 10 μM LY2157299 and 55 nM SB202190 (henceforth referred to as complete IntestiCult medium). Medium was changed every 2-3 days and were passaged after 7-10 days of culture. To passage, the medium was removed and the Matrigel containing enteroids was suspended in 1 mL of ice-cold DMEM/F12. The wells of enteroids were pooled and centrifuged at 400 x *g* for 5 minutes. The supernatant removed, the culture made to 1 mL suspension, and the enteroids were sheared by mechanical pipetting. The sheared enteroids were counted using a Brightfield microscope, and the fragments were suspended in Matrigel so that 200 fragmented enteroids were present in 50 μL of Matrigel per well. 650 μL fresh complete IntestiCult medium was used to culture the enteroids.

The enteroids were infected as described in (Blake et al., 2022). Briefly, the enteroids were pooled and sheared as described above. The number of enteroid fragments were counted and suspended in the appropriate volume of inoculum for each recombinant *E. coli* strain so that there was a final MOI of 100 per bovine cell. This is based on the calculation that each fragmented enteroid contained 100 cells (Blake et al., 2022).

The recombinant *E. coli* strains were prepared for infection as described previously, and the sheared enteroids were exposed to the *E. coli* inoculum in a suspension of complete IntestiCult medium. The enteroid and bacteria suspension were incubated at 37°C for 2 hours, pelleted at 500 rpm and washed with PBS. This was repeated twice more, and the enteroids lysed using 0.1% triton X-100.

The cell lysates were plated onto LB plates containing 100 μg/mL carbenicillin and 35 μg/mL chloramphenicol.

### Confocal microscopy

To image the infection of mammalian cell lines by recombinant *E. coli*, cells were cultured on coverslip slides prior to infection and fixed with 2% paraformaldehyde (PFA) for 20 minutes at room temperature. For the bovine enteroid samples, the samples were first pelleted at 400 x *g* and resuspended in 2% PFA (w/v) and fixed for 1 hour at 4°C. Samples to be stained were then permeabilised with 0.1% Triton-X 100 (v/v) for 15 minutes at room temperature and blocked with PBS containing 0.5% bovine serum albumin (v/v) and 0.02% sodium azide (w/v) (termed blocking buffer) for 30 minutes at room temperature. The samples were incubated with the designated primary antibody (Supplementary Table 1) diluted in blocking buffer for 1 hour at room temperature, washed three times with PBS, and incubated with the appropriate secondary antibody diluted in blocking buffer for 1 hour. Samples were then washed three times and nuclei stained with 300 nM DAPI for 2-5 minutes. The coverslips were mounted on a glass slide using Prolong Gold.

Enteroids were adhered to the surface of a glass slide using the CytoSpin method described in (Blake et al., 2022). Slides were visualised using Leica LSM710 upright immunofluorescence microscope.

## Results

### Expression of *mce* genes in MAP upon exposure to acidic conditions

Several adhesins expressed by MAP have been identified by quantifying gene upregulation in response to exposure to an acidic environment which may mimic that of the route of infection MAP experiences *in vivo*. To this end, the gene expression fold change of selected *mce* genes in MAP K10 and MAP C49 upon exposure to pH 3.0 7H9 medium for 2 hours was analysed using RT-qPCR using the geometric mean of *1g2* and *GAPDH* genes as internal controls. The change in *mce* gene expression was compared to *mce* gene expression in the corresponding MAP cultures which had been exposed to standard 7H9 medium for 2 hours. Both strains of MAP significantly upregulated *mce1A* and *mce1D* gene expression after exposure to acidic pH, whereas there was no change in *mce3C* and *mce4A* gene expression by either strain (Figure 1). This indicated *map1A* and *map1D* may therefore act as effector molecules which aid MAP infection of host cells. Similar studies have reported a resistance to acidic pH in MAP as a high CFU may still be recovered from treated cultures. In our hands, similar CFU/mL values were reported for MAP K10 post treatment as previous studies at 10^8^ CFU/mL, but this remains a significant reduction in viable bacteria compared to cultures which were not treated with acidic medium (Supplementary figure 1). MAP C49 seems to show a greater sensitivity to acidity, with the CFU/mL decreasing from 10^8^ CFU/mL to 10^7^ CFU/mL post treatment.

**Figure 1.**
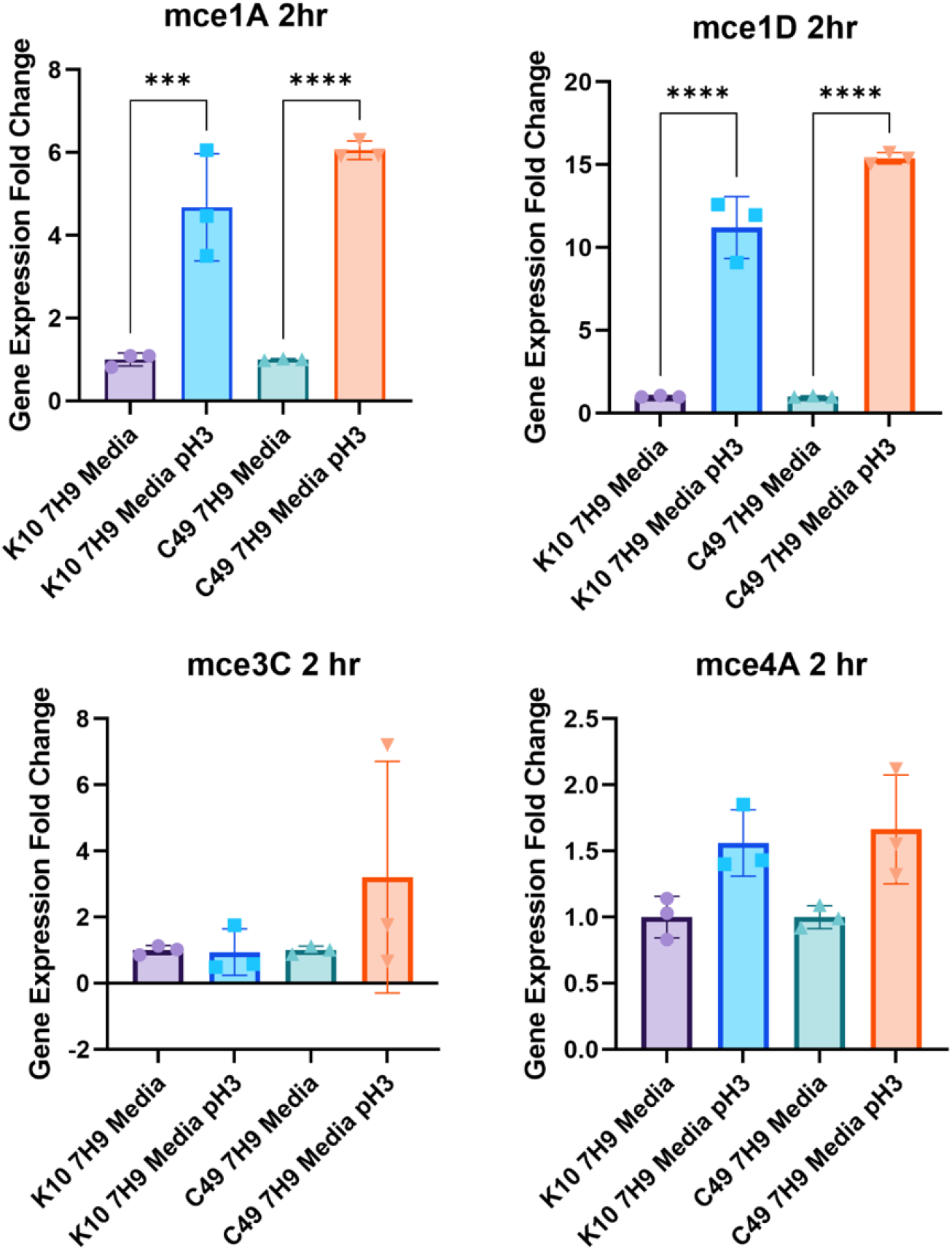
Regulation of MAP mce expression upon exposure to an acidic pH. The expression of mce genes was determined by RT-qPCR and calculated as fold change relative to the expression of gapdh and 1g2 as endogenous reference genes. Total RNA was isolated from 3 separate cultures of MAP K10 and C49 cultured to an OD_600_ 0.6 and pelleted to be resuspended in standard 7H9 media or pH3.0 7H9 media. Data presented as the mean of the fold change in gene expression from 3 separate cultures ±SD. Statistatical analysis performed using a 1-way ANOVA followed by a post hoc Dunnett’s test. P<0.05 = *; P<0.01 = **; P<0.001 = ***; P<0.0001 = ****.

### Cloning, expression and purification of Mce proteins

The expression of the His tagged Mce proteins was studied in *E. coli* Rosetta 2 (BL-21). The proteins were detected using anti-His-tag monoclonal antibody (Raybiotech) and were of the expected size, ranging from 43 – 60 kDa. The expression of the proteins was confirmed to be in the membranous fraction of the *E. coli* following IPTG induction using sub cellular fractionation of the bacteria (Figure 2). DNAK is a known cytoplasmic protein in *E. coli* and was used to ensure separation of the cytoplasmic and cell membrane fractions of the bacteria for Western Blotting. All Mce proteins were confirmed to be expressed in the cell membrane and absent from the cytoplasmic fraction, with the exception of Mce1D which was detected in both fractions of the *E. coli* (Figure 2).

**Figure 2.**
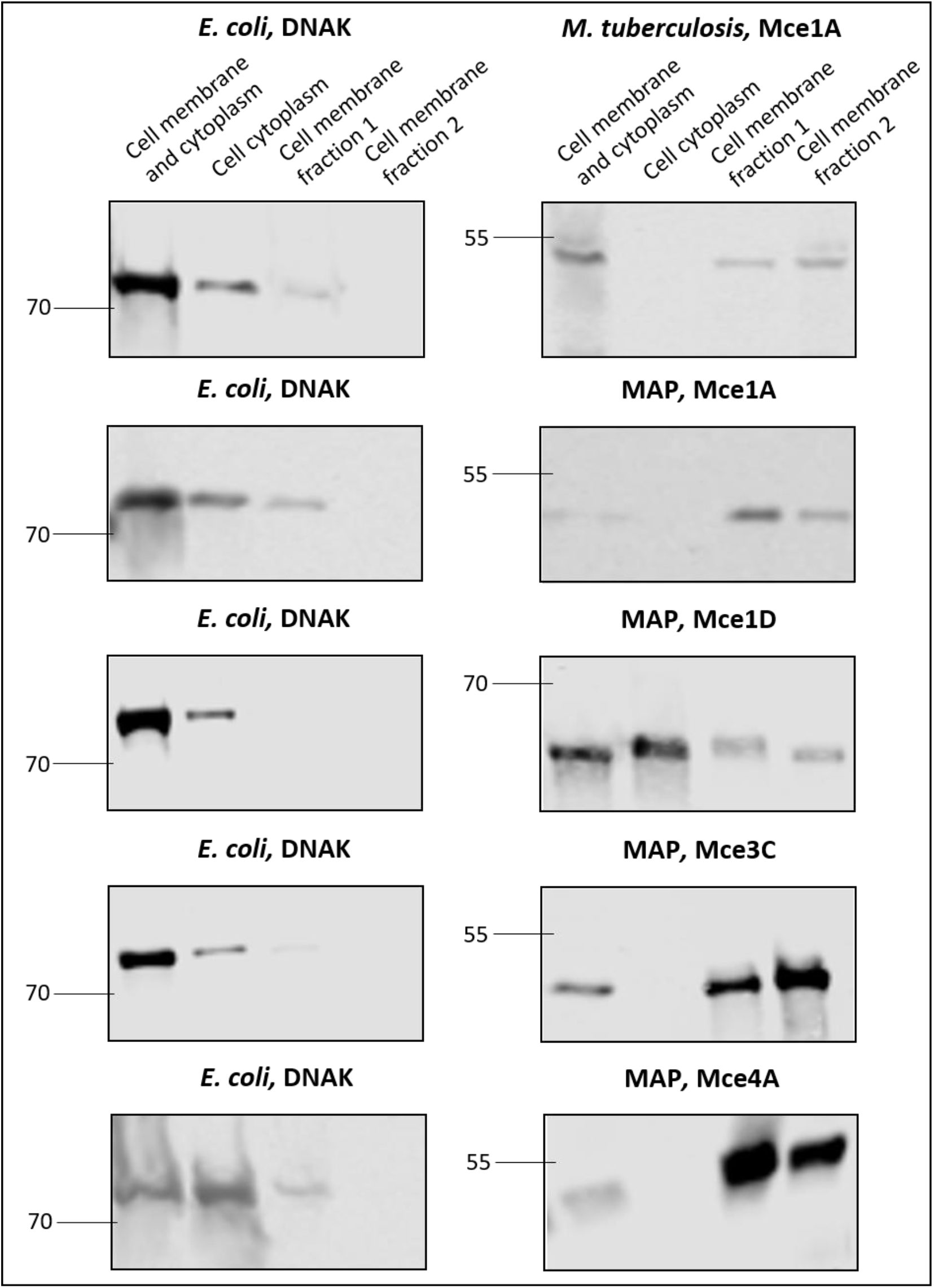
Western blot of E. coli strains expressing Mce protein after subcellular fractionation. E. coli clones were induced with 0.1 mM IPTG to produce their respective Mce protein for 2 hours at 37°C. The bacteria was then separated into fractions of the cell membrane and cytoplasm, the cytoplasm alone and 2 separate washes of the cell membrane. The fractions were separated by SDS-PAGE and electro-transferred to a nitrocellulose membrane. Rabbit monoclonal anti-His antibody was used to detect the His-tagged Mce protein to determine its location in the bacteria. Rabbit monoclonal anti-DNAK antibody was used as an E. coli cytoplasmic control. Numbers on the left lane indicate Molecular weight of protein standards in the ladder (kDa).

### Invasion of mammalian cells by recombinant *E. coli* expressing Mce protein

The use of *E. coli* as expression hosts of Mycobacterium derived Mce proteins was confirmed to have no significant effects on the viability of the bacteria compared to the empty plasmid control prior to performing invasion assays (Supplementary Figure 2). The invasion of the bovine epithelial cell line, MDBK cells, by recombinant *E. coli* was measured using CFU present in the cell lysate post infection. The results show significantly greater numbers of *E. coli* expressing Mce1A derived from *M tuberculosis* (Mtb1A) and *E. coli* expressing Mce1D derived from MAP (Map1D) were present in the cell lysate compared to the *E. coli* transformed with the empty pET21b(+) vector 2 hours post infection (Figure 3). This increase remained at 6 hours post infection for the *E. coli* expressing Mtb1A as the positive control, but not for the *E. coli* expressing Map1D. No other Mce protein aided the attachment or invasion of *E. coli* to bovine epithelial cells.

**Figure 3.**
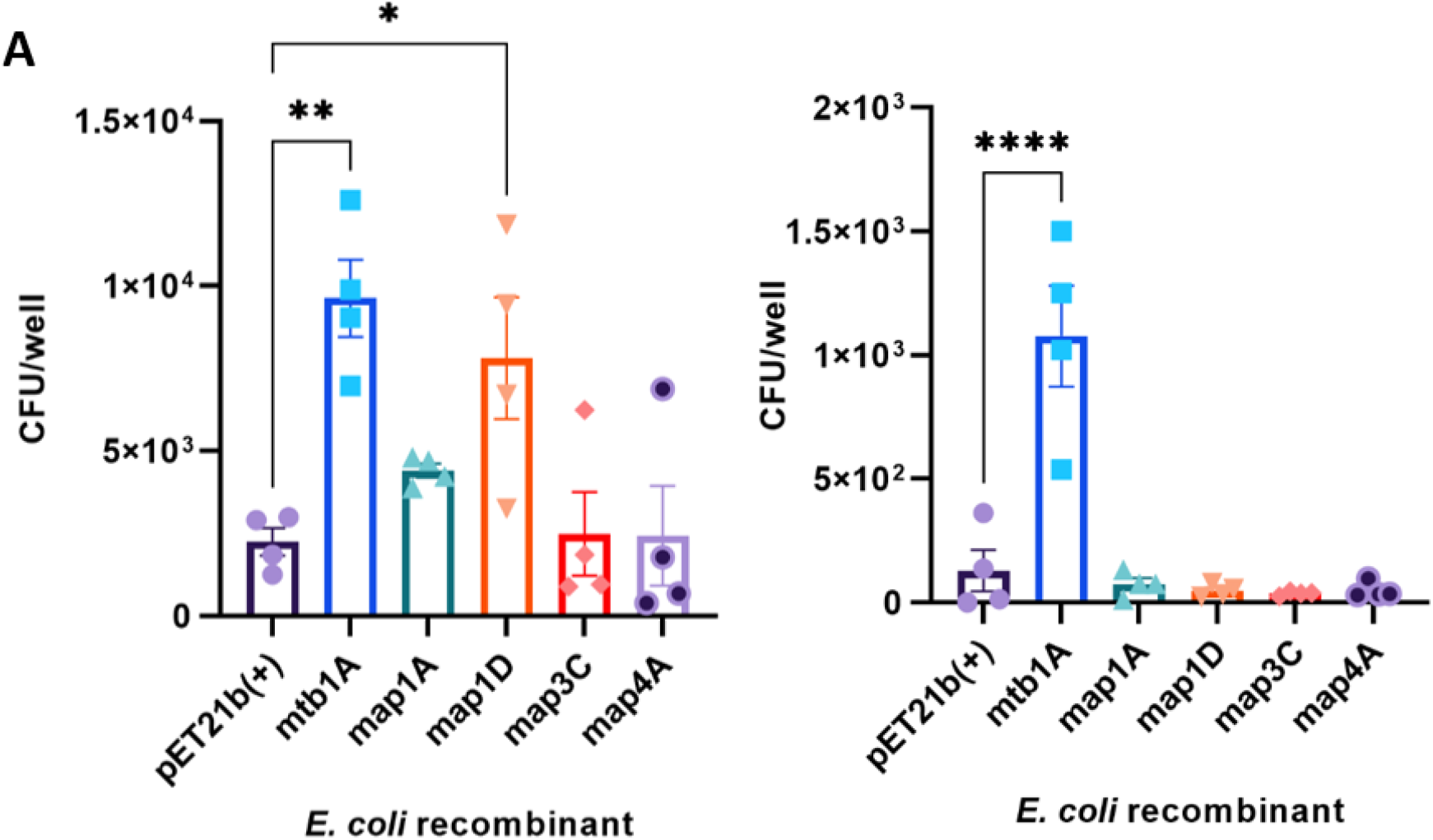
Attachment and survival of E. coli recombinants in MDBK cells. Mce protein expression was induced in E. coli recombinants with 0.1 mM IPTG for 2 hours at 37°C and used to infect MDBK cells at MOI 20. Cells were washed at 1 hour post infection and incubated for a further 1 or 5 hours. Cell lysates were plated for CFU analysis at 2 hours post infection **A)**; and at 6 hours post infection **B)**. Error bars presented as SEM of four biological replicates each performed with three technical repeats. Statistical analysis performed as a one-way ANOVA followed by a post hoc Dunnett’s test. P< 0.05 =*; P<0.001 =**; P<0.0001=***.

To investigate the function of the MAP derived Mce proteins in the context of a phagocytic cell line, the recombinant *E. coli* expressing Mce proteins were used to infect THP-1 cells. THP-1 cells have previously been used to investigate the function of Mce proteins using *E. coli* expression hosts, and may indicate the ability of these proteins to confer increased intracellular survival to the *E. coli*. After infection of the THP-1 cells for 2 and 6 hours by the *E. coli*, the cell lysates were plated to calculate the CFU/well. The results show significantly greater numbers of *E. coli* expressing Mtb1A and Mce1A derived from MAP (Map1A) were present in the cell lysate compared to the empty vector control (Figure 4). This difference persisted 6 hours post infection, although the total CFU/well decreased at 6 hours post infection compared to the CFU/well recovered 2 hours post infection (Figure 4).

**Figure 4.**
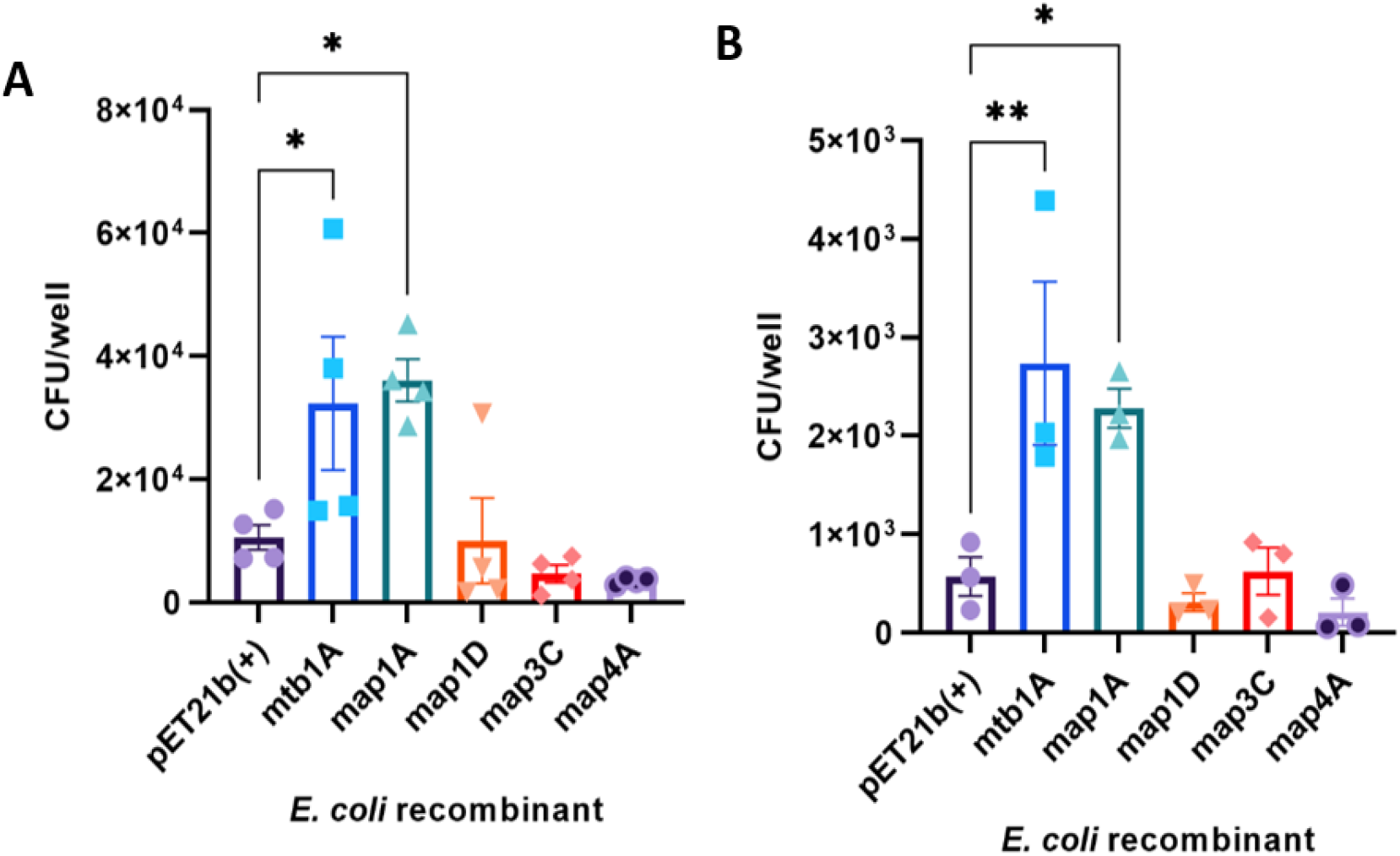
Uptake and survival of recombinant E. coli by THP-1 cells. Mce protein expression was induced in E. coli recombinants 0.1 mM IPTG for 2 hours at 37°C and used to infect THP-1 cells at MOI 10. Cells were incubated with media containing 10 μg/mL gentamicin after 1 hour infection and incubated for a further 1 or 5 hours. Cell lysates were plated for CFU analysis at 2 hours post infection (n=4) **A)**; and at 6 hours post infection (n=3) **B)**. Error bars presented as SEM of the specific number of biological replicates each performed with three technical repeats. Statistical analysis performed as a one-way ANOVA followed by a post hoc Dunnett’s test. P< 0.05 =*; P<0.001 =**; P<0.0001=***.

The invasion of MDBK and THP-1 cells by recombinant *E. coli* expressing Mtb1A, Map1A and Map1D was monitored by confocal microscopy. In THP-1 cells, all *E. coli* recombinants were observed to be intracellular at 2 and 6 hpi due to the phagocytic action of the mammalian cells. There were noticeably greater numbers of *E. coli* expressing Mtb1A or Map1A present in the cells at 6 hpi compared to the empty vector control (Figure 5). In MDBK cells at 2 hpi *E. coli* expressing Mtb1A and Map1D were bound to the surface of the of the MDBK cells, while there was no observable *E. coli* transformed with the empty pET21b(+) vector attached to the surface of the mammalian cell. At 6 hpi *E. coli* expressing Mtb1A were observed to be intracellular, whereas Map1D expressing *E. coli* remained on the surface of the cells in lower abundance compared to 2 hpi (Figure 5). There was no evidence of membrane ruffling for the *E. coli* located in the intracellular space, which may require EM to determine the mechanism of entry provided by the Mce protein being expressed.

**Figure 5.**
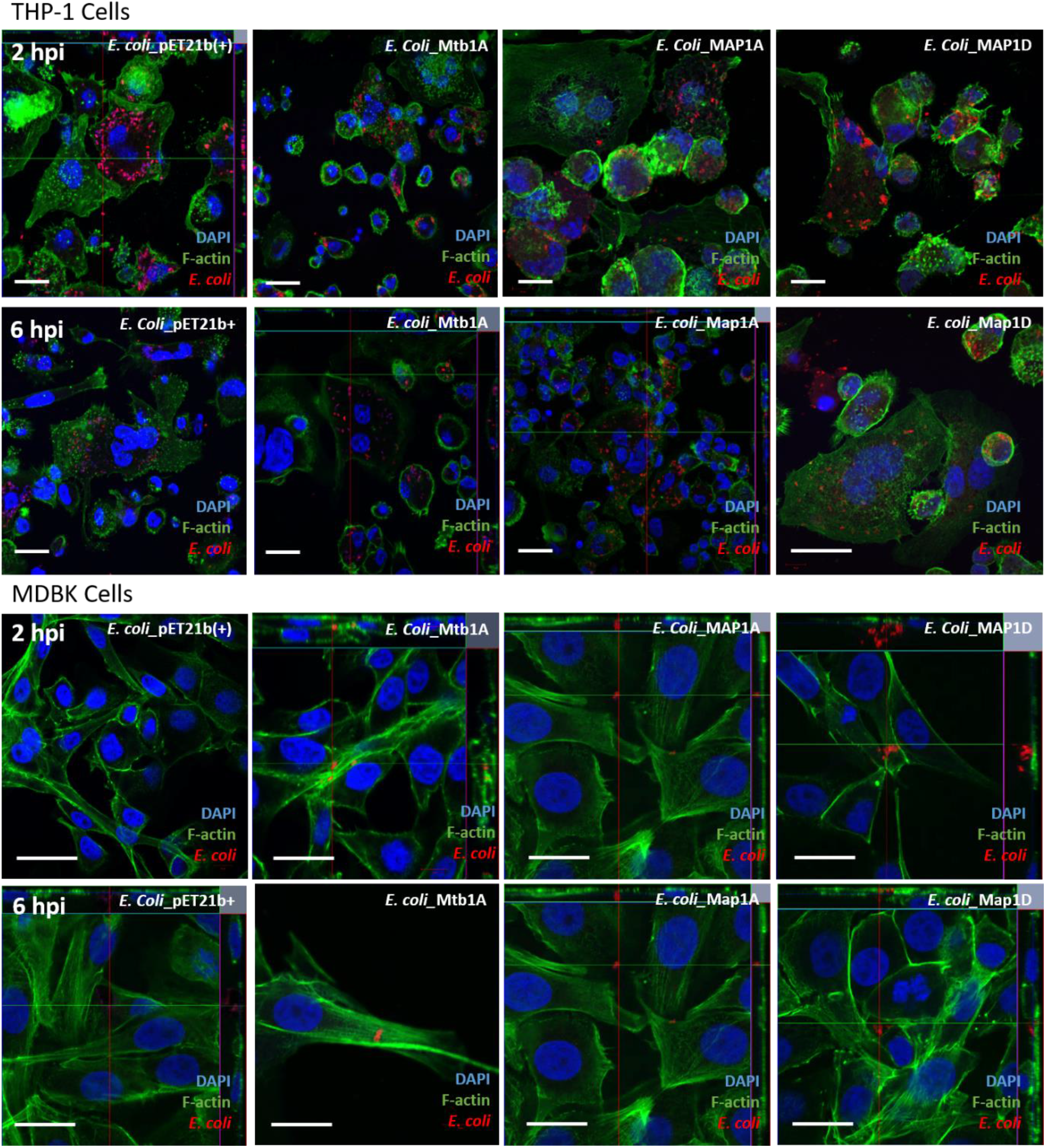
IF staining of infected mammalian cells with recombinant E. coli. The attachment and invasion of recombinant E. coli expressing Mce protein was visualised using IF staining and confocal microscopy at 2 and 6 hours post-infection. The cells were stained for nuclei (DAPI, blue), F-actin (Phalloidin, green) and anti-E. coli antibody (red). Scale bar = 20 μm.

### Invasion of bovine enteroids cells by recombinant *E. coli* expressing Mce protein

To further investigate the roles of Map1A and Map1D, invasion assays were performed in 3D basal-out bovine intestinal organoids (enteroids). These enteroids acted as a more physiologically representative model of the bovine intestine, where it is hypothesised the *mce* genes are involved in the attachment and invasion of the intestinal epithelial cells. Enteroids were sheared to expose the apical surface of the cells and incubated in suspension with recombinant *E. coli* expressing the relevant Mce protein for 2 hours at 37°C. The samples were lysed to calculate the CFU/well and the % inoculum was calculated for each recombinant *E. coli* investigated to account for differences in the inoculum between experiments (Figure 6). Interestingly, unlike the outcomes observed in the previous cell line studies, *E. coli* expressing both Map1A and Map1D are present in significantly greater numbers than the vector only control, pET21b(+), whereas Mtb1A did not confer an increased invasive capacity using the same model. This may indicate there is a species or organ specificity between these proteins despite their high homology at the protein level. The overall level of infection remains low at less than 2% inoculum, which is not unexpected due to the lack of phagocytic cells present in the enteroid cultures.

**Figure 6.**
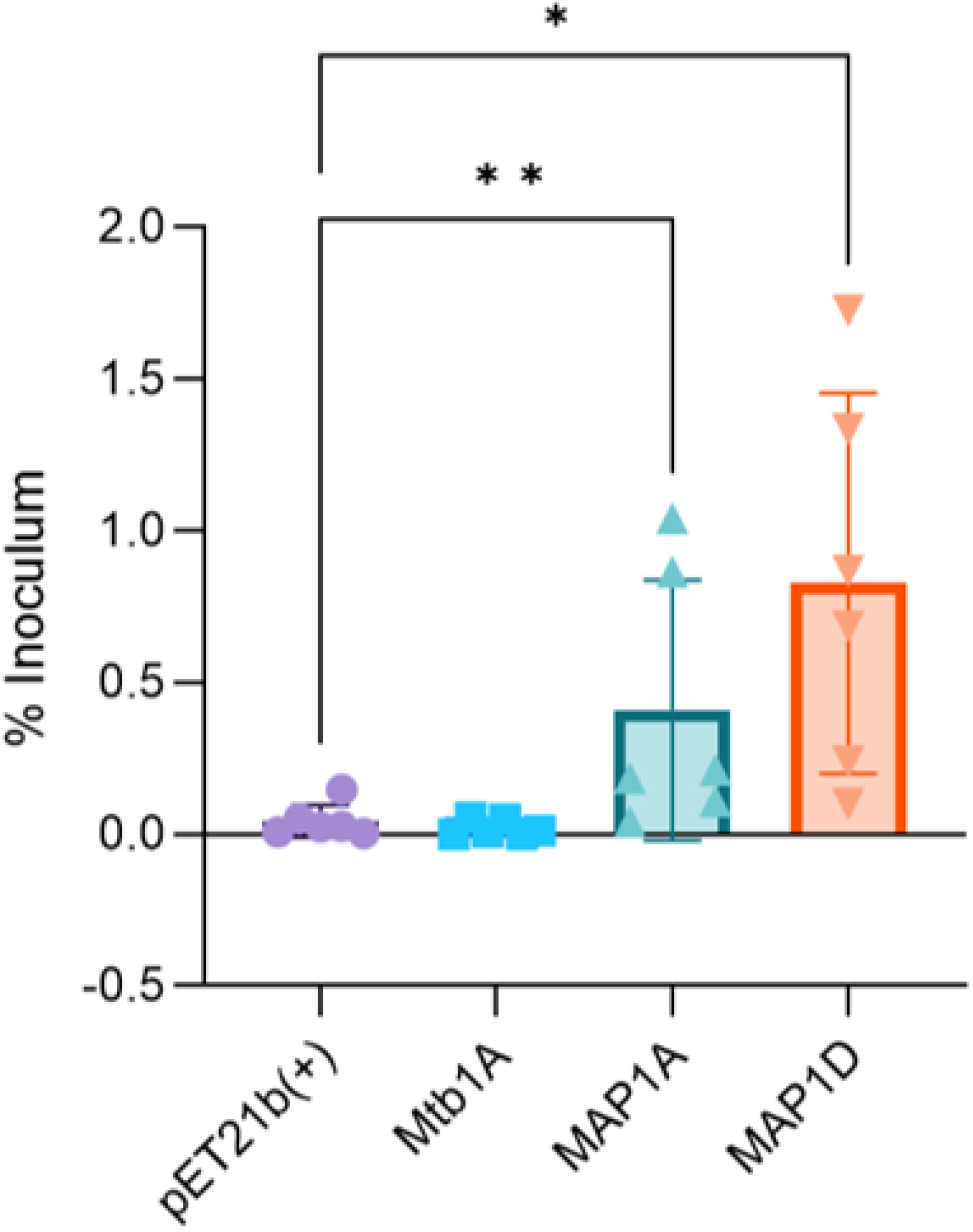
Infection of 3D basal-out bovine enteroids by E. coli recombinants. Mce protein expression was induced in E. coli recombinants with 0.1 mM IPTG for 2 hours at 37°C and used to infect enteroids in suspension at MOI 100 for 2 hours. Enteroids were washed three times with PBS. Cell lysates were plated for CFU quantification and data plotted as % of the inoculum. Data is representative of 6 biological replicates from enteroids derived from 2 separate calves. Error bars presented as SD. Statistical analysis performed as a one-tailed student’s T-test compared to the empty vector control. P< 0.05 =*; P<0.01=**.

## Discussion

The initial interaction between MAP and the host occurs at the intestinal lining and is a critical step in MAP pathogenesis which may impact the outcome of disease. Identification of effector molecules which play a role in this initial interaction increases our understanding of MAP pathogenesis and may provide novel therapeutic targets.

There are many studies that have identified MAP effector molecules by investigating gene regulation upon exposing MAP to conditions it would experience in a natural infection setting. Typically, exposure to milk and acidity has been observed to upregulate the expression of effector molecules which aid the attachment and invasion of host cells (Everman et al., 2018; Secott et al., 2001). From such experiments, fibronectin attachment proteins have been shown to be upregulated in response to acidic treatment (Secott et al., 2001), which then aids MAP attachment and invasion of M cells in the intestinal lining. However, these proteins were not observed to affect the ability of MAP to infect enterocytes which are known cell targets of the bacteria to aid translocation across the intestinal lining (Secott et al., 2004).

Proteins in the Mce family in pathogen Mycobacteria have been observed to aid the attachment and invasion of the bacteria to host epithelial cells. More recently, the *mce5* cluster has been identified as important for MAP virulence (Hemati et al., 2018), and mutation of *mce1D* resulted in a reduced capacity of MAP to infect MDBK cells (Alonso-Hearn et al., 2008).

In this study, a role for *mce1A, mce1D, mce3C* and *mce4A* in the initial interaction between MAP and the host was investigated. Consistent with other effector molecules, both *mce1A* and *mce1D* were upregulated in acidic conditions. Furthermore, three separate cell culture models were used to investigate the effect the expression of these Mce proteins would confer to their *E. coli* expression host. Both *mce1A* and *mce1D* were observed to increase attachment and invasion of either THP-1 or MDBK cells, but each *mce* gene did not infect both. Interestingly, both genes allowed an increase in the infection of 3D bovine enteroids when expressed by *E. coli*, whilst *mce1A* derived from *M. tuberculosis* did not. This indicates these genes exert a degree of host and cell type specificity, and that both cell types and receptors for the Mce1A and Mce1D MAP derived proteins are present in the bovine enteroid model. This lends credence to the hypothesis that these genes are involved in the initial interaction between MAP and the host at the intestinal lining.

While a role for *mce3C* and *mce4A* could not be elucidated in the current study, this does not definitively mean these genes do not act as effector molecules under different conditions.

Previously, Mce3C derived from *M. tuberculosis* was expressed on the surface of beads to monitor invasion of the beads in HeLa cells (Zhang et al., 2018). In our study, MAP derived *mce3C* was expressed in a Gram-negative *E. coli* host. It may be the outer membrane present in the Gramnegative *E. coli* interferes with the expression of Mce3C in the appropriate morphology on its surface.

In other Mycobacteria, Mce4A was found to be upregulated in bacteria in the stationary phase of growth (Kumar et al., 2003). This stage is likened to what a bacterium may experience later in the infection process once becoming intracellular. Therefore, this proteins may not be involved in the initial interaction, but it may be more influential in longer infection studies which incorporate cholesterol uptake to maintain MAP viability. In this study, only the initial interaction was studied within the first 6 hours, and this may not be optimal to understand the role of Mce4A.

Furthermore, it is only through the use of the enteroids as multicellular model which allowed the observation of the invasive phenotypes of both *mce1A* and *mce1D* in a single model. This indicates that the model chosen to investigate function of MAP genes should be carefully considered in future experiments as cell lines do not reflect the complexity of the *in vivo* host which MAP infects.

## Acknowledgements

We would like to acknowledge our colleagues for their comments and suggestions for experimental design and the Biotechnology and Biological Sciences Research Council Institute Strategic Programme Grant (BBS/E/D/20002173) for funding this work.

## Declarations

The authors declare that the research was conducted in the absence of any commercial or financial relationships that could be construed as a potential conflict of interest.

## Supplementary

**Supplementary Table 1.** A table outlining the antibodies used in this paper and the corresponding concentration. *Indicates antibodies used for Western Blots.

| Antibody/staining agent | Concentration | Manufacturer Number |
| --- | --- | --- |
| Anti-His* | 1µg/mL | RB-10-0002-100 |
| Anti-Dnak* | 1µg/mL | A207645 |
| Anti-E. coli | 10 µg/mL | ab137967 |
| Anti-Rabbit IgG H+L* | 1µg/mL | 06/2019 |
| Anti-Mouse IgG H+L* | 1 µg/mL | 06/2016 |
| Phalloidin 488 | 66 µM | A12379 |
| Anti-Rabbit 594 | 10 µg/mL | Z25307 |

**Supplementary Figure 1.**
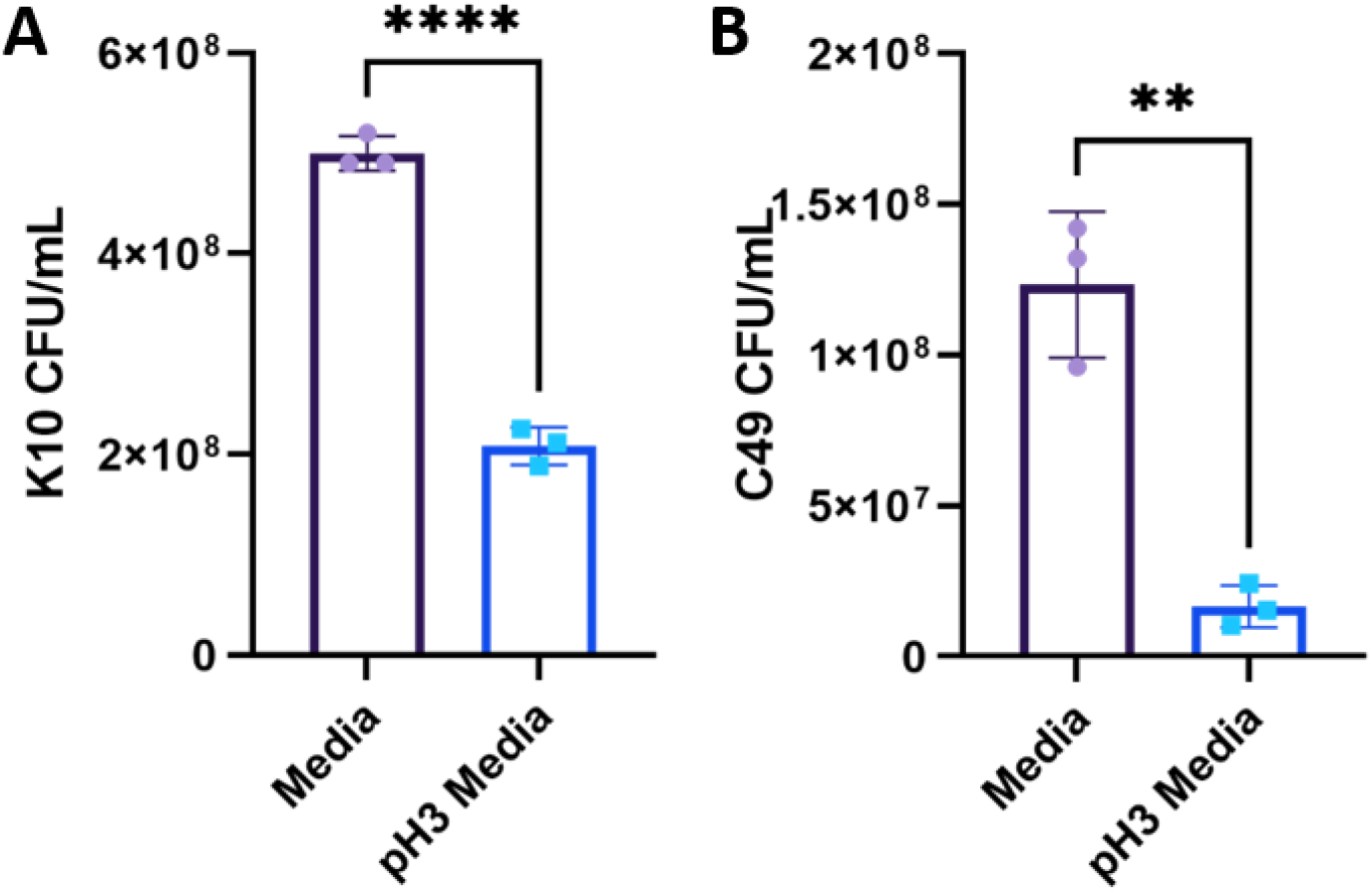
CFU/ml values of MAP cultured in acidic 7H9 media. MAP was cultured to an OD_600_ 0.6 and pelleted. The pellet was re-suspended in 7H9 growth media that was either the standard pH or pH 3.0 and cultured at 37°C 100 rpm for 2 hours. The cultures were then diluted and plated onto 7H10 agar and incubated at 37°C for up to 6 weeks. A) MAP K10 CFU values; B) MAP C49 CFU values. Data analysed using Student’s unpaired T-test. P<0.05 = *; P<0.01 = **; P<0.001 = ***; P<0.0001 = ****.

**Supplementary Figure 2.**
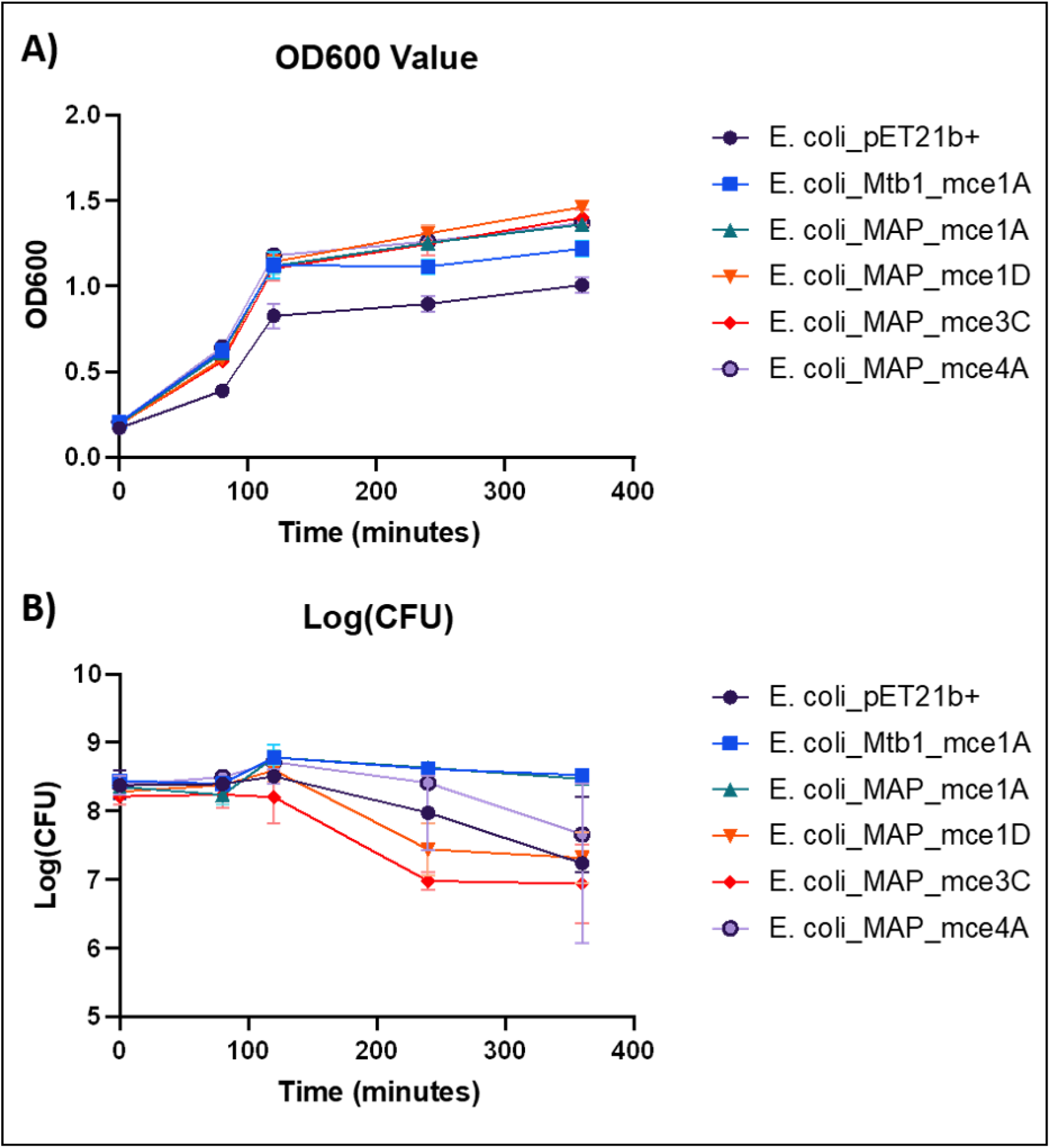
Growth curves of recombinant E. coli mutants upon induction of Mce protein expression. 1:10 dilution was performed from an overnight culture of recombinant E. coli clones and cultured at 37°C. OD_600_ **A)** and CFU/mL **B)** values were taken at the indicated times, and protein production was induced with 0.1 mM IPTG upon an OD_600_ value of 0.6 being reached. Bacteria were cultured on LB agar containing the relevant antibiotics and cultured overnight at 37°C for CFU/mL analysis. Results were gained from 3 biological replicates.

## References

Alonso-Hearn, M., Patel, D., Danelishvili, L., Meunier-Goddik, L., & Bermudez, L. (2008). The Mycobacterium avium subsp. paratuberculosis MAP3464 gene encodes an oxidoreductase involved in invasion of bovine epithelial cells through the activation of host cell Cdc42. Infection and Immunity, 76(1), 170–178. https://doi.org/10.1128/IAI.01913-06

Arruda, S., Bomfim, G., Knights, R., Huima-Byron, T., & Riley, L. W. (1993). Cloning of an M. tuberculosis DNA fragment associated with entry and survival inside cells. Science, 261(5127), 1454–1457. https://doi.org/10.1126/science.8367727

Arsenault, R. J., Maattanen, P., Daigle, J., Potter, A., Griebel, P., & Napper, S. (2014). From mouth to macrophage: mechanisms of innate immune subversion by Mycobacterium avium subsp. paratuberculosis. 45, 1–15. https://doi.org/10.1186/1297-9716-45-54

Bannantine, J., Huntley, J., Miltner, E., Stabel, J., & Bermudez, L. (2003). The Mycobacterium avium subsp. paratuberculosis 35 kDa protein plays a role in invasion of bovine epithelial cells. Microbiology (Reading, England), 149(Pt 8), 2061–2069. https://doi.org/10.1099/MIC.0.26323-0

Bermudez, L. E., Petrofsky, M., Sommer, S., & Barletta, R. G. (2010). Peyer’s Patch-Deficient Mice Demonstrate That Mycobacterium avium subsp. paratuberculosis Translocates across the Mucosal Barrier via both M Cells and Enterocytes but Has Inefficient Dissemination. Infection and Immunity, 78(8), 3570. https://doi.org/10.1128/IAI.01411-09

Casali, N., & Riley, L. W. (2007). A phylogenomic analysis of the Actinomycetales mce operons. BMC Genomics, 8, 60. https://doi.org/10.1186/1471-2164-8-60

Chitale, S., Ehrt, S., Kawamura, I., Fujimura, T., Shimono, N., Anand, N., Lu, S., Cohen-Gould, L., & Riley, L. W. (2001). Recombinant Mycobacterium tuberculosis protein associated with mammalian cell entry. Cellular Microbiology, 3(4), 247–254. https://doi.org/10.1046/j.1462-5822.2001.00110.x

Corn, J. L., Manning, E. J. B., Sreevatsan, S., & Fischer, J. R. (2005). Isolation of Mycobacterium avium subsp. paratuberculosis from free-ranging birds and mammals on livestock premises. Applied and Environmental Microbiology, 71(11), 6963–6967. https://doi.org/10.1128/AEM.71.11.6963-6967.2005

Everman, J. L., Danelishvili, L., Flores, L. G., & Bermudez, L. E. (2018). MAP1203 Promotes Mycobacterium avium Subspecies paratuberculosis Binding and Invasion to Bovine Epithelial Cells. Frontiers in Cellular and Infection Microbiology, 8(JUN), 217. https://doi.org/10.3389/FCIMB.2018.00217

Hemati, Z., Haghkhah, M., Derakhshandeh, A., Singh, S. V., & Chaubey, K. K. (2018). Cloning and characterization of MAP2191 gene, a mammalian cell entry antigen of Mycobacterium avium subspecies paratuberculosis. Molecular Biology Research Communications, 7(4), 165–172. https://doi.org/10.22099/mbrc.2018.30979.1354

Koressaar, T., & Remm, M. (2007). Enhancements and modifications of primer design program Primer3. Bioinformatics, 23(10), 1289–1291. https://doi.org/10.1093/BIOINFORMATICS/BTM091

Kumar, A., Bose, M., & Brahmachari, V. (2003). Analysis of Expression Profile of Mammalian Cell Entry (mce) Operons of Mycobacterium tuberculosis. Infection and Immunity, 71(10), 6083. https://doi.org/10.1128/IAI.71.10.6083-6087.2003

Larsen, A. B., Merkal, R. S., & Cutlip, R. C. (1975). Age of cattle as related to resistance to infection with Mycobacterium paratuberculosis. American Journal of Veterinary Research, 36(3), 255–257.

Lombard, J. E. (2011). Epidemiology and Economics of Paratuberculosis. Veterinary Clinics of North America: Food Animal Practice, 27(3), 525–535. https://doi.org/10.1016/J.CVFA.2011.07.012

Nielsen, S. S., & Toft, N. (2008). Ante mortem diagnosis of paratuberculosis: A review of accuracies of ELISA, interferon-γ assay and faecal culture techniques. In Veterinary Microbiology (Vol. 129, Issues 3–4, pp. 217–235). Elsevier. https://doi.org/10.1016/j.vetmic.2007.12.011

Pfaffl, M. (2001). A new mathematical model for relative quantification in real-time RT-PCR. Nucleic Acids Research, 29(9), E45. https://doi.org/10.1093/NAR/29.9.E45

Schleig, P. M., Buergelt, C. D., Davis, J. K., Williams, E., Monif, G. R. G., & Davidson, M. K. (2005). Attachment of Mycobacterium avium subspecies paratuberculosis to bovine intestinal organ cultures: Method development and strain differences. Veterinary Microbiology, 108(3–4), 271–279. https://doi.org/10.1016/j.vetmic.2005.04.022

Secott, T. E., Lin, T. L., & Wu, C. C. (2001). Fibronectin attachment protein homologue mediates fibronectin binding by Mycobacterium avium subsp. paratuberculosis. Infection and Immunity, 69(4), 2075–2082. https://doi.org/10.1128/IAI.69.4.2075-2082.2001

Secott, T. E., Lin, T. L., & Wu, C. C. (2004). Mycobacterium avium subsp. paratuberculosis fibronectin attachment protein facilitates M-cell targeting and invasion through a fibronectin bridge with host integrins. Infection and Immunity, 72(7), 3724–3732. https://doi.org/10.1128/IAI.72.7.3724-3732.2004

Untergasser, A., Cutcutache, I., Koressaar, T., Ye, J., Faircloth, B. C., Remm, M., & Rozen, S. G. (2012). Primer3—new capabilities and interfaces. Nucleic Acids Research, 40(15), e115–e115. https://doi.org/10.1093/NAR/GKS596

Wang, M., Gao, Z., Zhang, Z., Pan, L., & Zhang, Y. (2015). Human Vaccines & Immunotherapeutics Roles of M cells in infection and mucosal vaccines Roles of M cells in infection and mucosal vaccines. https://doi.org/10.4161/hv.36174

Zhang, Y., Li, J., Li, B., Wang, J., & Liu, C. H. (2018). Mycobacterium tuberculosis Mce3C promotes mycobacteria entry into macrophages through activation of β2 integrin-mediated signalling pathway. Cellular Microbiology, 20(2). https://doi.org/10.1111/CMI.12800

